# Egg viability decreases rapidly with time since ovulation in the rainbow darter: implications for the costs of choosiness

**DOI:** 10.1101/199430

**Authors:** Rachel L. Moran, Rachel M. Soukup, Muchu Zhou, Rebecca C. Fuller

## Abstract

Egg viability in the rainbow darter *Etheostoma caeruleum*, a fish apparently lacking female mate choice, was found to decline rapidly after ovulation. We observed that the majority of a female’s clutch may fail to hatch if she is prevented from mating for as little as six hours. These data suggest that exercising female mate preferences may be selectively disfavoured in *E. caeruleum* due to the high cost of delaying mating.

---

The degree of discrimination exhibited by female animals when choosing a mate ranges from none to extreme. This diversity exists because female mate choice is shaped by a number of variables, including the subset of males made available to her by environmental factors and male-male interactions (Beehler & Foster, 1988; Jennions & Petrie, 1997; Wong & Candolin, 2005), and the costs of mate choice behaviour (Janetos, 1980; Real, 1990; Kokko *et al.*, 2003). If the costs of choosing outweigh the benefits, the most favoured strategy would be to mate with the first male the female encounters, i.e. random mating. Several costs to female mate choice (e.g. time and energy expenditure, predator exposure) have been investigated across a variety of taxa (reviewed in Reynolds & Gross, 1990; Jennions & Petrie, 1997), but one has thus far received little attention: decline in gametic quality over time. In fishes, the phenomenon of decreasing egg viability post-ovulation, termed egg overripening, has been reported across a range of species including Atlantic salmon *Salmo salar* Linnaeus 1758 (de Gaudemar & Beall, 1998), goldfish *Carassius auratus* (Linnaeus 1758) (Formacion *et al*., 1993), Atlantic halibut *Hippoglossus hippoglossus* (Linnaeus 1758) (Bromage *et al*., 1994), and turbot *Scophthalmus maximus* (Linnaeus 1758) (McEvoy, 1984). Having eggs susceptible to overripening would pressure females to spawn quickly, as waiting could incur a substantial fitness cost. Such selection for rapid mate acquisition may consequently lead to a decrease in female choosiness and/or influence the expression of female mating preferences.

The rainbow darter *Etheostoma caeruleum* Storer 1845 is a small benthic fish common in freshwater streams across the eastern United States (Page, 1983). During the breeding season from late March to early June, brightly coloured males attempt to guard gravid females from rival males. Females signal their readiness to spawn by performing nosedigs, wherein she pusheher head into the gravel. Spawning involves the female burying herself shallowly in gravel with an arched posture; once a male takes position above her, both vibrate and release eggs and sperm. No parental care is practiced (Winn, 1958; Fuller, 2003).

Although there is evidence that female *E. caeruleum* favour some types of males in dichotomous trials, such preferences have little apparent effect on the outcome of mating due to male-male competition (Fuller, 2003). In any case, female *E. caeruleum* demonstrate no overt choice when allowed to interact freely with males: having assumed the spawning position, the female always spawns with the first male to arrive regardless of his characteristics (pers obs), suggesting that exercising choice may be disadvantageous. Good reasons exist to suspect that mate choice in female *E. caeruleum* may harbour a cost in prolonging the time between ovulation and spawning: in previous experiments, females have been observed expelling and subsequently eating unfertilized eggs when held in isolation (Zhou & Fuller, 2014). Furthermore, isolated females subsequently allowed to spawn with males often produce entirely inviable clutches (R. Moran, pers obs). This study aimed to formally test the hypothesis that delaying spawning is costly for female *E. caeruleum*, by quantifying change in egg viability as a function of time since ovulation.

*E. caeruleum* were collected by kick seine from Mill Pond Outlet (Kalamazoo Co., Michigan) in April and May 1998 (year 1) and from an unnamed tributary of the Saline Branch Drainage Ditch (Champaign Co., Illinois) in March 2017 (year 2). Fish from year 1 were maintained at the Kellogg Biological Station. Fish from year 2 were maintained at the University of Illinois at Urbana-Champaign. In both years, fish were housed in male-female pairs in 38 litre aquariums with gravel substrate, maintained at external ambient temperature and light:dark cycle. The fish were fed frozen bloodworms (chironomid larvae) and live tubifex worms twice per day.

Fish were monitored during daylight hours over four to five days post-capture. A female was assumed to have recently ovulated when she performed a nosedig. After a female performed a nosedig, she was moved to an empty tank. Females were kept in isolation for various lengths of time: 0 hours (n=10), 6 hours (n=6), 12 hours (n=7), 24 hours (n=5). Following the isolation period, a male was introduced and the fish were allowed to freely spawn. The resulting eggs were collected with a siphon and placed in small tubs filled with water; dilute methylene blue was added to inhibit fungal growth. Hatching success was recorded as the number of eggs yielding fry out of total number of eggs collected. To test for a relationship between hatching success and female isolation time, we used a quasibinomial regression with a logit link function. To determine whether there was an effect of collection year and location on hatching success, year was included as a covariate in the model. The quasibinomial error distribution was used to account for overdispersion in the response variable (i.e., hatching success). Statistical analysis was performed in R (version 3.4.0).

All females spawned following the introduction of a male. The number of eggs collected ranged from 17 to 110, and was uncorrelated with isolation time or year. Hatching success declined strongly as a function of increasing female isolation time (F_1,25_ = 5.91, p < 0.05; Fig. 1). There was no effect of year on hatching success (F_1,25_ = 3.30, p = 0.08). Although hatching success varied at each female holding time (Fig. 1), these data suggest that on average, greater than 50% of a female’s clutch is likely to become non-viable if retained for as little as six hours after ovulation.

**Figure 1.**
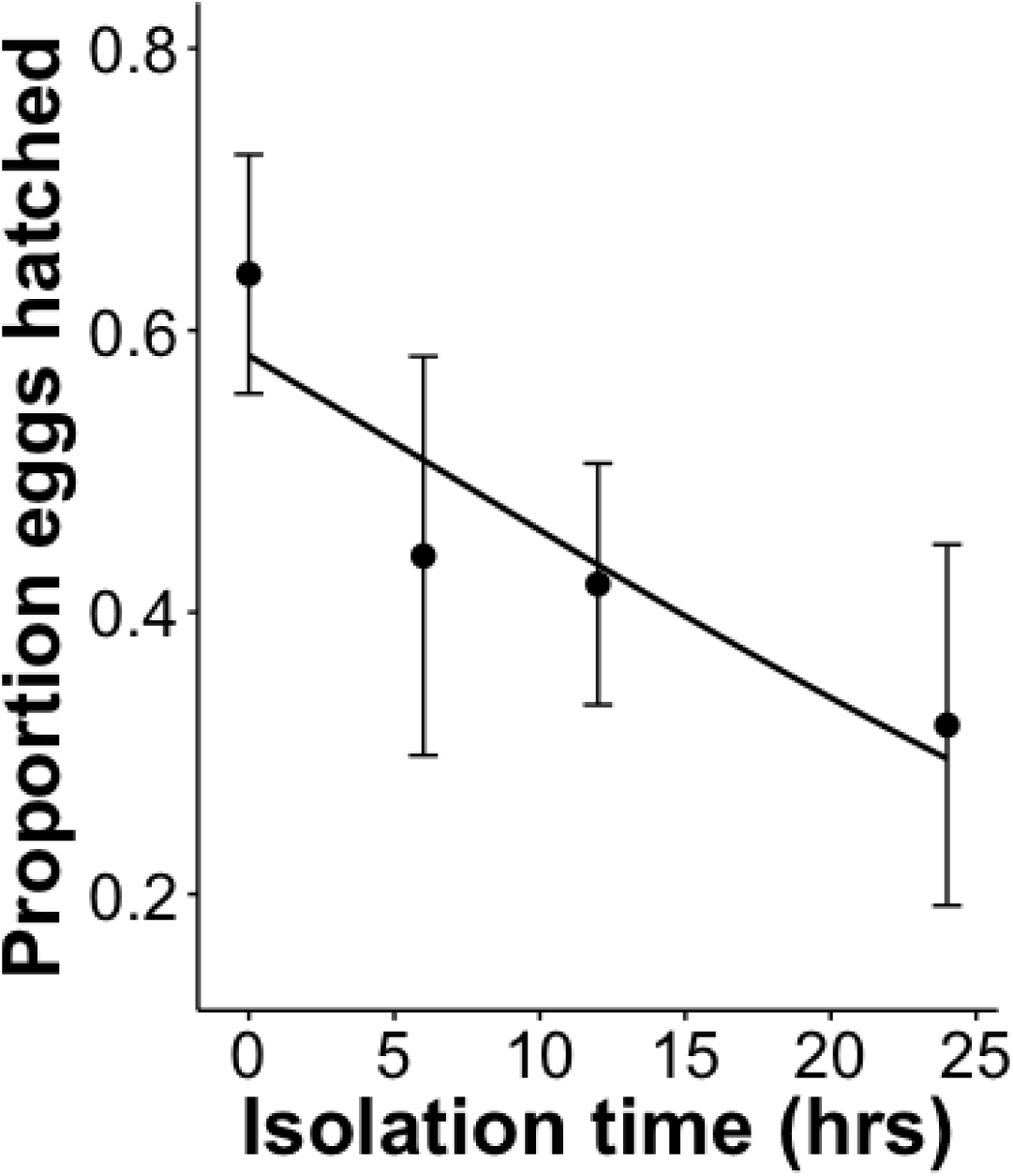
The proportion of eggs that hatched from a clutch decreased with increasing time that a female was held in isolation (i.e. prevented from spawning) after ovulation occurred.

Given that hatching success declines precipitously over time, gravid female *E. caeruleum* appear to be under strong pressure to fully spawn a clutch of eggs in less than 24 hours. Female who fail to do so risk substantial egg mortality. The time constraint for spawning may be exacerbated by the fact that female *E. caeruleum* release only a small fraction of ovulated eggs per spawning bout and thus must spawn multiple times to fully expel an entire clutch (Fuller, 1998). Under these conditions, the cost for a female *E. caeruleum* to reject a male may be unacceptably high.

Female mate choice in *E. caeruleum* may be further disfavoured by strong male-male competition. Male *E. caeruleum* fight vigorously for access to gravid females, attempting to monopolize spawning and prevent the participation of “sneaky” males (Winn, 1958; Fuller, 2003). Hence, the choice of males within a single patch is likely limited to those that are competitively superior. Furthermore, there may be little additional benefit for the female to choose following male-male competition if male competitive ability predicts fitness benefits to females and her offspring (Wong & Candolin, 2005).

Darters are a highly speciose clade that have received increasing attention from evolutionary biologists over the past decade. Spectacular and diverse male colouration in darters has been suggested to act as an agent of speciation by sexual selection, with the most commonly posited mechanism being divergent female mate choice (Mendelson, 2003; Williams & Mendelson, 2010; Williams *et al*., 2013). However, evidence is mounting that female choice is limited in atleast some darter species (Pyron, 1995; Fuller, 2003; Zhou *et al*. 2015; Moran *et al.*, 2013, 2017). This study suggests that female preference in darters is likely costly due to the need to spawn shortly after ovulation while egg viability remains high. Considering that egg overripening seems to be common across a variety of fish species, often occurring over time frames comparable to or shorter than in *E. caeruleum* (Kjørsvik *et al*., 1990), its importance as an evolutionary force on mate choice may be underappreciated.

## Acknowledgements

This work was supported by the Cooperative State Research, Education, and Extension Service, US Department of Agriculture, under project number ILLU 875-952, the National Science Foundation (DEB 0953716 and IOS 1701676), and the University of Illinois. The treatment of animals was approved by the Institutional Animal Care and Use Committee under protocol #17031.

